# Dissipation of lysosome pH impairs formation and collapses existing LPS-induced lysosome tubules in macrophages

**DOI:** 10.1101/2025.09.25.677195

**Authors:** Shiraz Anwar, Solmaz Ebrahimi-Iranpour, Maria R. Narciso, Jennifer Li, Saaimatul Huq, Tracy Lackraj, Tamara Beriia, Nazia Chowdhury, Mojca Mattiazzi Usaj, Roberto J. Botelho

## Abstract

While lysosomes are typically globular in morphology, lipopolysaccharide-activated macrophages reorganize lysosomes into an expanded tubular network. Here, we sought to determine if the V-ATPase and the lysosomal pH contribute to LPS-mediated lysosome tubulation in macrophages. We found that inhibition of the V-ATPase prevented lysosome tubulation and collapsed preformed tubules. However, the V-ATPase also controls mTORC1 activity, which itself is needed for tubulation. To distinguish between lysosomal pH and mTORC1, we turned to NH_4_Cl, which alkalinized lysosomal pH but did not interfere with mTORC1; yet NH_4_Cl blocked tubulation showing that an acidic lysosomal pH is needed for lysosome remodelling. Moreover, clamping the pH to either acidic or alkaline pH caused tubules to collapse, indicating that the pH gradient promotes tubulation, rather than a specific pH. These effects were not due to altered microtubule organization or impaired lysosome motility, suggesting that motors remained associated with lysosomes. On the other hand, while LPS did not alter the average pH of spherical or tubular lysosomes, growing tubules displayed a more acidic peripheral end relative to the pericentral end. Based on this observation, we propose that a localized pH gradient along the tubule may enable tubulation by modulating factors that catalyse tubule growth.

**Significance statement:** Macrophages and dendritic cells exposed to lipopolysaccharides convert their globular lysosomes into a tubular lysosome network. This requires microtubules, kinesin and dynein motors, and mTORC1 activity.

We show that the lysosome pH gradient is required to form and maintain tubular lysosomes in macrophages. While global lysosomal pH did not change in response to LPS and tubulation, we observed that growing tubules typically had a more acidic peripheral end, which may promote tubulation. We propose that lysosomal pH gradient directly or indirectly coordinates tubulation machinery, possibly coordinating motor activity to deform the lysosomes into a tubule.

## Introduction

Lysosomes are the terminal point of endocytosis, phagocytosis, and autophagy, receiving a wide range of molecular, particulate, and organellar cargo for degradation and component recycling. Thus, lysosomes are ideally positioned to integrate the nutrient and environmental conditions of cells to elicit a cellular adaptation to these conditions (Luzio *et al*., 2007; Settembre *et al*., 2013; Raben and Puertollano, 2016; Inpanathan and Botelho, 2019). To achieve these functions, lysosomes are equipped with >60 hydrolases and numerous membrane transporters to recycle amino acids and divers metabolites, ion channels to modulate ion flow, and the V-ATPase, which acidifies the lumen. These properties and functions are enabled by a network of regulatory factors like the Rab7 and Arl8b GTPases, lipid signals such as phosphoinositides, kinases like mTORC1, motor proteins, and fusion and fission complexes (Luzio *et al*., 2007; Schwake *et al*., 2013; Inpanathan and Botelho, 2019; Saffi and Botelho, 2019; Ballabio and Bonifacino, 2020).

The acidic milieu of lysosomes is often described as ideal for hydrolytic activity. Nonetheless, the H^+^ gradient is also important for other functions including driving the co-transport of metabolites, the flow of counterions such as Cl^-^, Ca^2+^, and Na^+^, and recruitment of specific proteins to the lysosomal membrane (Christensen *et al*., 2002; Mundy *et al*., 2012; Colacurcio and Nixon, 2016; Löbel *et al*., 2022; Zhang *et al*., 2022; Wu *et al*., 2023). In addition, the lysosomal H^+^ gradient likely regulates trafficking proteins. For example, acidification of endosomes and late endosomes helps recruit Arf1/COPI and the Arf6/ARNO complexes to their membrane (Aniento *et al*., 1996; Hurtado-Lorenzo *et al*., 2006). Additionally, we showed that acidification of endosomes and phagosomes helps terminate phosphatidylinositol-3-phosphate signaling (Naufer *et al*., 2018). Thus, the function of the lysosomal H^+^ gradient goes beyond providing an optimal milieu for degradation.

In most cells, lysosomes are a collection of functionally heterogeneous globular organelles. For example, peripheral lysosomes are more alkaline than perinuclear lysosomes (Johnson *et al*., 2016). Moreover, lysosomes can undergo significant change upon specific triggers. This is most obvious in lipopolysaccharide (LPS)-exposed macrophages and dendritic cells, which convert their globular lysosomal system into a tubular network within two hours of exposure (Hipolito *et al*., 2018, 2019). In addition, these cells expand the total lysosomal volume to accommodate higher levels of fluid phase uptake (Swanson *et al*., 1987a; Hipolito *et al*., 2019). This is driven by mTOR and enhanced translational activity (Hipolito *et al*., 2019). In turn, lysosome remodelling is associated with improved antigen presentation and transport of muramyl dipeptides across lysosomal membranes (Chow *et al*., 2002; Nakamura *et al*., 2014; Hipolito *et al*., 2019). Lysosome tubulation itself is dependent on Rab7 and Arl8b GTPases and their effectors RILP, FYCO, and SKIP1, which engage dynein and kinesin to elongate lysosomes on microtubule tracks (Mrakovic *et al*., 2012). It is not known how these antagonistic motors are coordinated to productively deform lysosomes into elongated tubules. One possibility is that H^+^ and Ca^2+^ gradients generate local gradient domains to modulate dynein and kinesin (Hipolito *et al*., 2018). In fact, pH and Ca^2+^-gradients were observed along the long axis of a subset of lysosome tubules induced by Tudor, a DNA nanodevice that binds to Ku70/80, and LPS in RAW macrophages (Suresh *et al*., 2021).

Here, we set out to further characterize the role of lysosomal pH in lysosome tubulation in LPS-treated primary macrophages. We observed that disrupting lysosomal pH with V-ATPase inhibitors or using weak buffers like NH_4_Cl prevented lysosome tubulation and collapsed an existing tubular network. This was not due to altered microtubule organization or inhibition of global lysosome motility activity. Instead, we observed that growing but not collapsing tubules were more acidic at their anterior tip. We propose that this environment helps coordinate motor activity to form lysosome tubules.

## Results

### V-ATPase inhibition blocks lysosome tubulation

The pH of lysosomes is central for lysosome function (Mindell, 2012; Xiong and Zhu, 2016). We thus hypothesized that lysosomal pH and the V-ATPase play a role in LPS-mediated lysosome tubulation in primary macrophages. To test this hypothesis, we pre-labelled lysosomes in macrophages with pHrodo-dextran and Alexa647-dextran and calculated their ratio; pHrodo fluorescence increases with lower pH, while the pH-independent Alexa647 was used to correct for differences in endocytosis and trafficking to lysosomes. This was followed by exposure to either vehicle or 10 µM concanamycin A (ConA) for 30 min to block the V-ATPase and then cells were stimulated for 2 h with LPS before live-cell imaging. First, we demonstrated that acute inhibition of the V-ATPase with ConA readily depleted the lysosomal pH in both resting and LPS-treated macrophages as indicated by the drop in the fluorescence intensity ratio of pHrodo-dextran to Alexa655-dextran (Sup. Fig. S1). Second, we manually binned macrophages in multiples of five lysosome tubules per macrophage (Fig. 1A, B) and determined the cumulative frequency of cells with more than 15 tubules (Fig. 1C). While most cells had <10 lysosome tubules in vehicle-treated cells, LPS significantly increased the number of cells with >15 tubules (Fig. 1A, C). Importantly, V-ATPase inhibition with ConA significantly declined basal and LPS-induced lysosome tubulation (Fig. 1A, C). We also tested whether the V-ATPase activity was necessary for lysosome tubule maintenance by preforming lysosome tubules with LPS and then exposing cells to ConA for 30 min after tubulation. We observed a collapse of the lysosome tubular network into globular lysosomes upon V-ATPase blockage (Fig. 2). Collectively, the V-ATPase activity is important for formation and maintenance of lysosomal tubules in LPS-treated macrophages.

**Figure 1:**
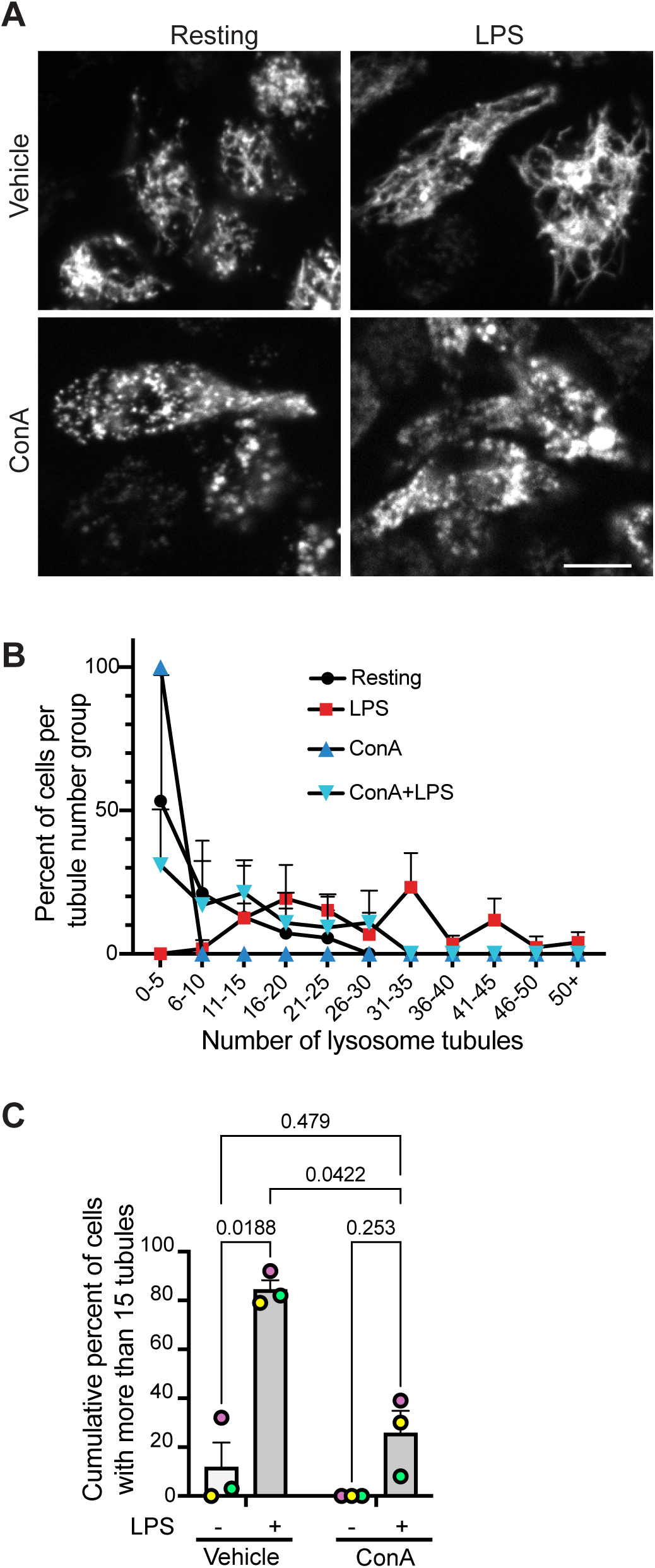
Inhibition of V-ATPase with ConA arrests LPS-mediated lysosome tubulation. **A.** Micrographs of primary macrophages showing lysosomes labelled with Alexa647-conjugated dextran. Cells were either treated with vehicle or with 1 μM concanamycin A (ConA) for 30 min followed by additional vehicle or 500 ng/mL of LPS for 2 h. Scale bar = 5 μm **B.** Percent frequency of cells binned into multiples of 5 lysosome tubules from three independent experiments using at least 100 cells per condition per experiment. Lines connecting data points are for presentation and do not indicate data. **C.** Cumulative percent cells with >15 tubules from data in B. Data are shown as mean ± SEM and were analysed by repeated measures two-way ANOVA and Tukey’s post-hoc test; p values are shown.

**Figure 2:**
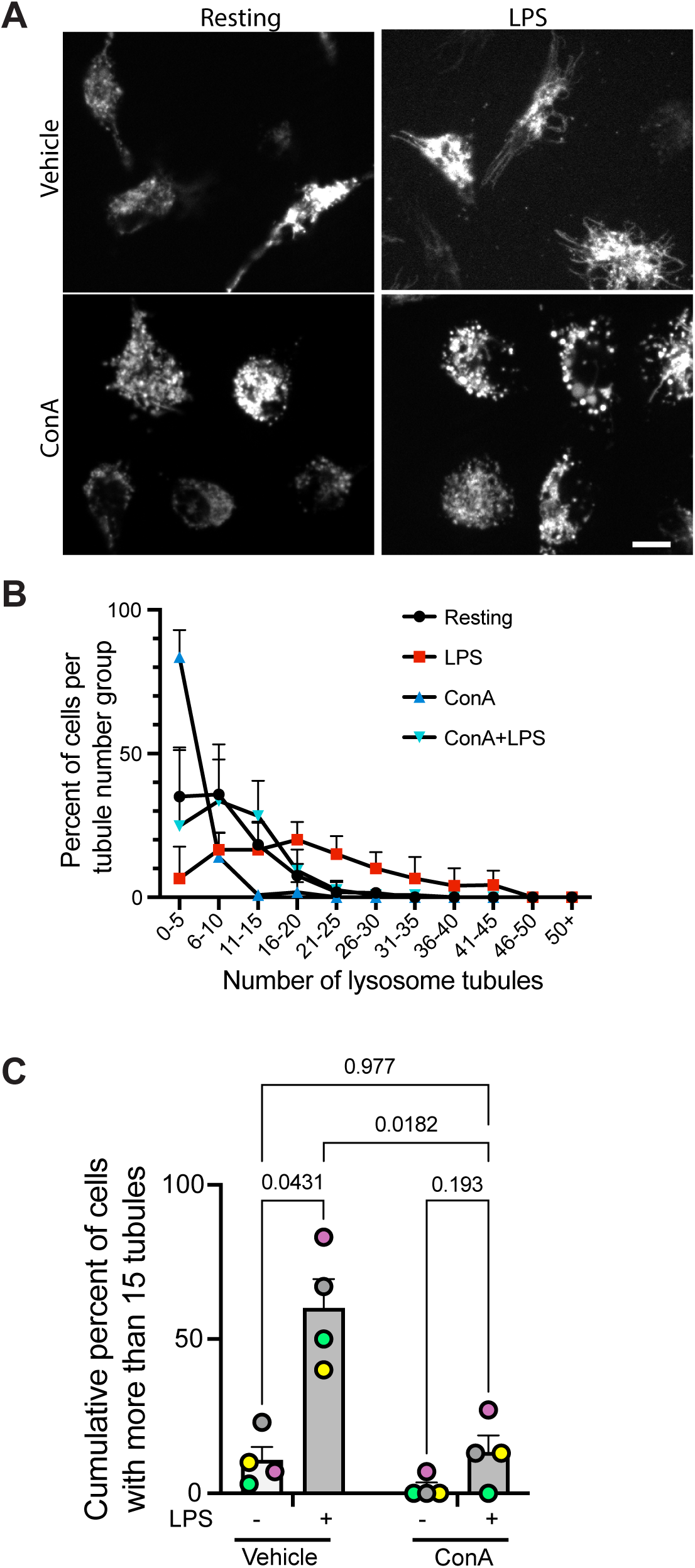
Inhibition of V-ATPase with ConA collapses pre-existing lysosome tubules. **A.** Micrographs of primary macrophages showing lysosomes decorated with Alexa647-dextran. Cells were first exposed to vehicle or LPS for 2 h to form tubules and then further treated for 30 min with vehicle or 1 µM ConA. Scale bar = 5 μm **B.** Percent frequency of cells binned into multiples of 5 lysosome tubules from three independent experiments using at least 150 cells per condition per experiment. Lines connecting data points are for presentation and do not indicate data. **C.** Cumulative percent cells with >15 tubules from data in B. Data are shown as mean ± SEM and were analysed by repeated measures two-way ANOVA and Tukey’s post-hoc test; p values are shown.

### V-ATPase inhibition, but not alkalinizing agents, blocks mTORC1

Our data show that V-ATPase activity is important for formation and maintenance of lysosome tubules in LPS-treated macrophages. Nevertheless, the V-ATPase also affects mTORC1 activity, which we previously revealed promotes lysosome expansion and tubulation (Zhang *et al*., 2014; Saric *et al*., 2016). To validate whether this is the case in macrophages, we tested the effect of ConA on mTORC1 activity by measuring the levels of phospho-S6K. As we previously observed, LPS promoted pS6K levels (Fig. 3A, B; (Saric *et al*., 2016; Hipolito *et al*., 2019)). In comparison, application of ConA abated this LPS-driven increase in pS6K levels (Fig. 3A, B). Thus, ConA-mediated inhibition of lysosome tubulation may be because of dissipated lysosomal pH gradient and/or inhibition of mTORC1.

**Figure 3:**
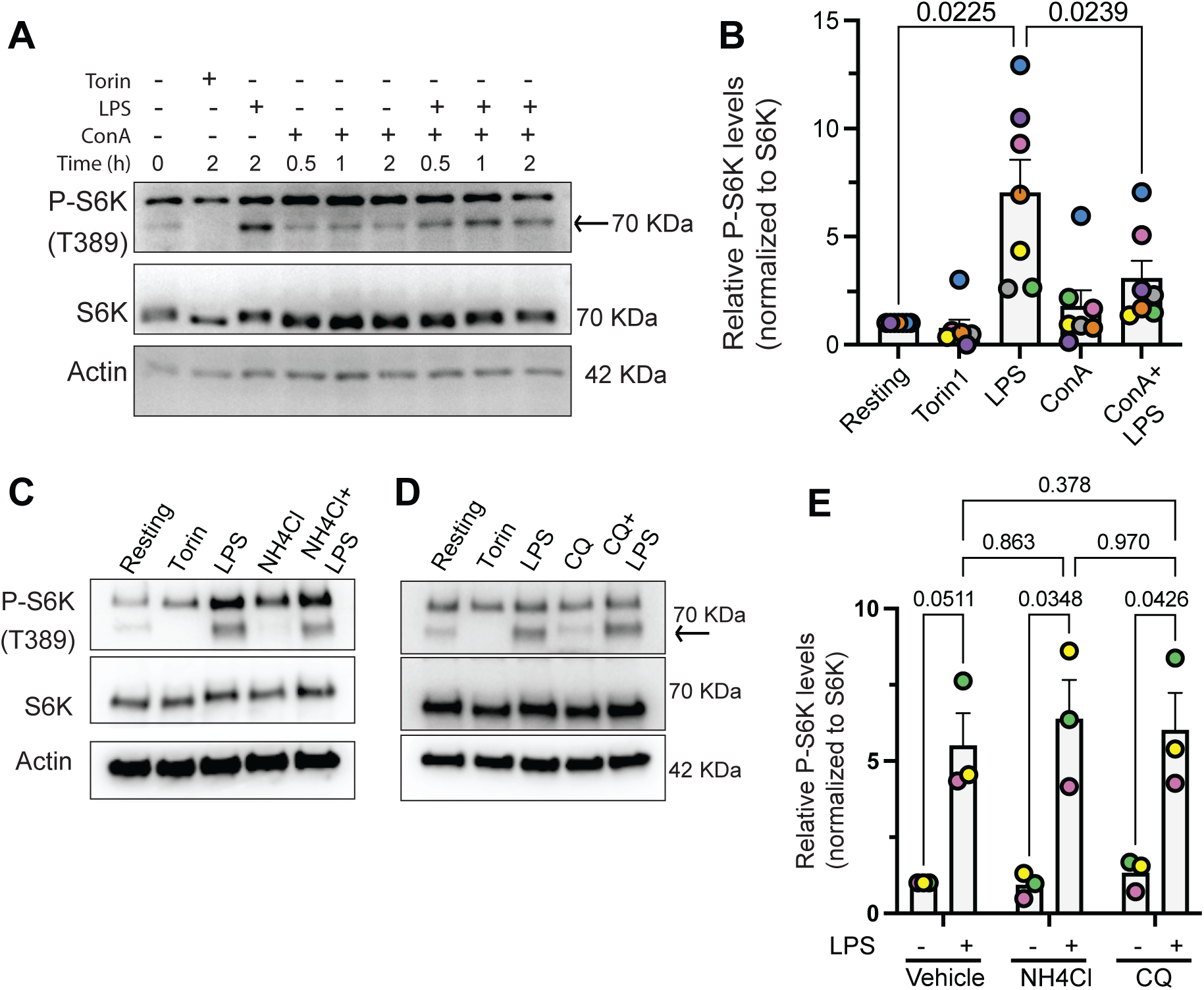
mTORC1 activity depends on the V-ATPase but not pH. **A, C** and **D.** Western blots of bone-marrow derived macrophages in resting state, or LPS-treated with and without ConA (A), NH_4_Cl (C), or chloroquine (CQ; D) and probed for β-actin, total S6 kinase (S6K), and p-T389 S6K (indicated by arrow). Torin-1 was used as a positive control for mTORC1 inhibition**. B** and **E.** Quantification of p-S6K signal corrected against total S6K for each condition and normalized to resting cells. Data are mean ± STD from n=7 (B) or n=3 (E) and analysed by repeated-measures one-way ANOVA and Dunnett’s post-hoc test (B) or two-way ANOVA and Tukey’s post-hoc test (E). Relevant p-values are shown.

To differentiate between these two possibilities, we turned to treatments expected to deplete proton gradients, but not the V-ATPase activity itself. We used the FCCP proton decoupler (Benz and McLaughlin, 1983), the NH_4_Cl weak base (Poole and Ohkuma, 1981; Christensen *et al*., 2002), and chloroquine, a putative alkalinizing agent (Poole and Ohkuma, 1981). First, FCCP treatment caused macrophage cell death within 2 h, thus we did not pursue this agent further (not shown). Second, we set out to confirm that NH_4_Cl and chloroquine could de-acidify lysosomes while not affecting mTORC1. We observed that NH_4_Cl and chloroquine did not affect basal or LPS-stimulated pS6K levels, unlike ConA (Fig. 3C-E). To ensure that lysosomes were alkalinized under both treatments, we measured the relative lysosomal pH as a ratio of pHrodo-dextran to Alexa647-dextran loaded in lysosomes. While NH_4_Cl deacidified lysosomes, we were surprised that chloroquine did not alkalinize the lysosomes despite extensive lysosome vacuolation whether measured by microscopy (Sup. Fig. S2A, B) or by flow cytometry (Sup. Fig. S2C). To exclude the possibility that this was dye-dependent, we measured the fluorescence ratio of Oregon green-to Alexa546-conjugated dextran by flow cytometry, whereby Oregon green fluorescence is quenched at more acidic pH – again, while NH_4_Cl appears to alkalinize lysosomes, this was not observed for chloroquine, which may in fact hyperacidify lysosomes (Fig. S2D). Importantly, others have observed that chloroquine does not alkalinize the pH of lysosomes despite clearly vacuolating these organelles (Yoon *et al*., 2010; Lu *et al*., 2017; Mauthe *et al*., 2018). Overall, our data suggest that NH_4_Cl can deplete lysosomal pH while not affecting mTORC1.

### Machine learning model to quantify lysosome tubulation

Before investigating the effects of NH_4_Cl (and chloroquine) in lysosome tubulation, we decided to test if machine learning could be used to score tubulation. Manual counting of lysosome tubules is labour intensive and sometimes difficult to discern where one tubule begins or ends. To address this, we implemented a transfer-learning approach by fine-tuning a ResNet50 convolutional neural network (CNN) using Keras and TensorFlow libraries within the Jupyter Notebook platform. We generated a training set containing at least 300 images of individual macrophages each manually annotated as either non-tubulating or tubulating (Sup. Fig. S3A shows examples of images for each category). After training, we evaluated the performance of this model on an independent set of macrophages, from those with no or few tubular lysosomes to those with high levels of lysosome tubulation. The model produced a binary output (tubulating vs non-tubulating). With the initial training set, the model correctly classified more than 80% of new images according to their manually assigned label (Sup. Fig. S3B, C). We then extracted the probability score assigned by the model for each image reflecting the likelihood of the cell belonging to the “tubulating group”, enabling statistical analyses between different treatments.

### Weak bases prevent lysosome tubulation in macrophages

To test if weak buffers impaired lysosome tubulation induced by LPS, we pre-loaded macrophages with Alexa647-dextran and then treated cells with NH_4_Cl or chloroquine for 30 min, followed by vehicle or LPS for 2 h. As before, cells treated with LPS exhibited extensive lysosome tubulation, while control cells had few (Fig. 4A, B). Importantly, administration of NH_4_Cl or chloroquine strongly impaired lysosome tubulation in LPS-treated cells (Fig. 4A, B). In fact, chloroquine tended to cause extensive lysosome vacuolation as previously observed (Fig. 4A, Fig. S2A). Whereas Fig. 4B shows manual scoring, we then used the machine learning model trained above. Single cells were cropped and categorized by treatment (Fig. 4C, shows examples) and then analysed by the trained lysosome tubulation model. In agreement with our visual inspection, resting cells had low average probability of being “cells with lysosome tubules”, while LPS-treated cells had a much higher probability (Fig. 4D). Treatment of macrophages with either chloroquine or NH_4_Cl with or without LPS all netted low probability scores of being “cells with lysosome tubules” (Fig. 4D). Thus, NH_4_Cl and chloroquine both disallowed lysosome tubulation, but likely via different mechanisms: NH_4_Cl dissipates lysosomal pH without affecting mTORC1 activity, suggesting that lysosomal pH is necessary for tubulation; chloroquine retains an acidic lysosomal pH, but causes massive swelling of lysosomes that may obstruct tubulation.

**Figure 4:**
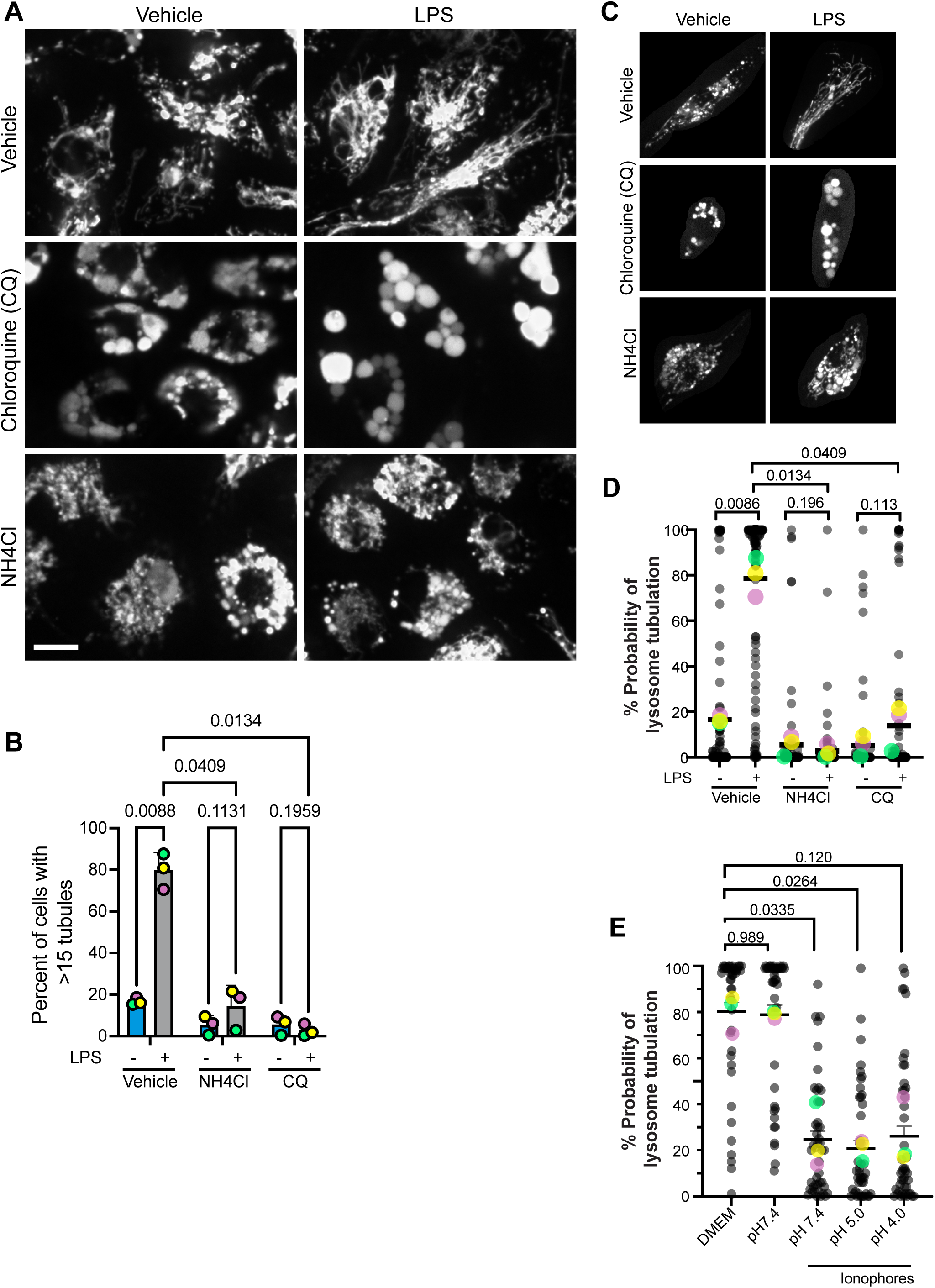
Weak bases disturb LPS-induced lysosome tubulation. **A.** Confocal single-plane images of living primary macrophages pre-labelled with Alexa647-dextran, treated or not with LPS, and/or 10 mM NH_4_Cl or 60 µM chloroquine (CQ). Scale bar = 10 µm. **B.** Percent cells with >15 lysosome tubules. **C.** Representative single-cell images from each condition cropped from original micrographs fed into the Jupyter machine-learning algorithm for lysosome tubulation analysis. **D.** Probability of a cell belonging to the “lysosome tubulation” category using our machine-learning model for cells treated with and without LPS and with or without indicated weak bases. **E**. Probability of a cell belonging to the “lysosome tubulation” category after pre-incubation with LPS to form tubules and then exposure to ionophores to clamp the cytoplasmic pH to pH 7.4, 5.0, or 4.0. For B, D, and E, data are shown as mean ± SEM from different independent experiments. At least 50 cells per condition per experiment were scored. For D and E, individual cell probability scores are shown as black dots. For B and D, data were analysed with a repeated measures two-away ANOVA and for E, data was tested with a one-way ANOVA. Tukey’s post-hoc test was applied to each; p values are indicated.

Next, we wondered if the lysosomal pH gradient or a specific pH value was required for lysosome tubulation. We addressed this question by using the monensin and nigericin ionophores to clamp the internal pH of cells to pH 4, 5, or 7.4 using macrophages pre-exposed to LPS to form tubules and then quantified the stability of lysosome tubules. Using our machine-learning model, lysosome tubules disintegrated in cells clamped to any of the tested pH values, though this did not meet statistical significance for pH 4 (Fig. 4E). We interpret these data to mean that the lysosomal pH gradient is essential for tubulation rather than a specific pH value.

### Lysosome tubules are more acidic at their growing tips

We next measured the lysosomal pH to try to understand how it sustains LPS-mediated lysosome remodelling. As described earlier, we co-loaded cells with pHrodo-dextran and Alexa647-dextran and calculated their ratio. This approach showed ConA-treated or NH_4_Cl-treated cells had more alkaline lysosomes than resting cells (Sup. Figs. S1 and S2). Interestingly, we did not observe a significant change in the cellular pHrodo/Alexa647 fluorescence ratio between resting and LPS-treated cells by microscopy (Fig. 5A, B) or by flow cytometry (Sup. Fig. S2C), indicating that global lysosomal pH does not change with LPS under the observed conditions. Moreover, segregating globular from tubular lysosomes did not reveal a difference in pH in either resting or in LPS-treated cells (Fig. 5A, C), consistent with prior observations (Suresh *et al*., 2021). We repeated these experiments with Oregon green-dextran normalized to Alexa546-conjugated dextran observing the same patterns by flow cytometry (Sup. Fig. S2D) and microscopy (Sup. Fig. S4A-C), whereby Oregon green fluorescence is quenched in acidic milieu. Collectively, we have no evidence that 2 h LPS exposure changes the lysosomal pH, though this may depend on time and level of exposure (Tsang *et al*., 2000; Baker *et al*., 2015).

**Figure 5:**
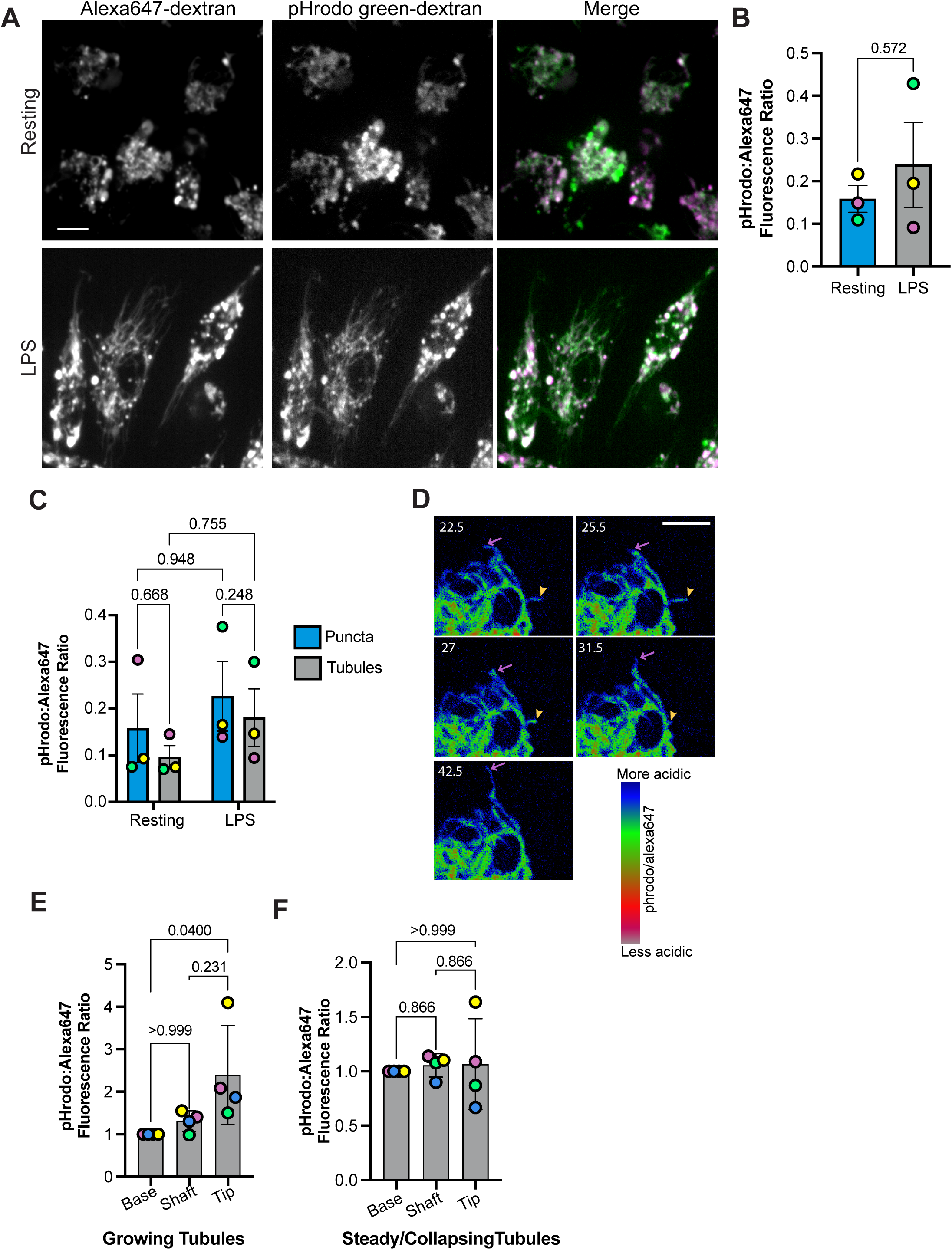
LPS does not change the global lysosomal pH, but growing tubules are more acidic at their anterior tips. **A.** Confocal images of resting and LPS-treated primary macrophages labelled co-labelled with pHrodo green dextran (pH sensitive) and Alexa647-dextran (pH insensitive); merge shows two channels (green = pHrodo; magenta = Alexa647). Scale bar = 10 µm. **B**. Average cellular pHRodo to Alexa647 fluorescence intensity in resting and LPS-treated macrophages. **C**. Mean pHRodo to Alexa647 fluorescence intensity of punctate and tubular lysosomes in resting and LPS-treated macrophages. **D.** Timelapse of ratiometric images of pHRodo/Alexa647 fluorescence of LPS-treated macrophages. The ratiometric images are pseudocoloured corresponding to pH shown in the LUT. The purple arrow identifies a tubular lysosome peripheral tip growing and showing more acidic ratio. The yellow arrow identifies a tubular lysosome collapsing inward and losing the more acidic ratio at the tip. Time is in seconds. **E, F.** Normalized mean ratio of pHRodo/Alexa647 fluorescence along the long axis of tubules observed to grow (E) or unchanged/collapsing (F); regions along the tubule axis were defined as the base (beginning of tubule at the perinuclear region), shaft (centre of the tubule), and tip (tubule end towards the cell periphery). All data are means ± SEM from at least three independent experiments. Data in B were from n=50 cells per experiment per condition and were tested by a paired, two-tail Student’s t-test; Data in C were based on n=25 cells and 1800-2000 individual lysosomes per experiment per condition and was tested with a repeated measures two-way ANOVA and Tukey’s post-hoc test. Data in E and F were assessed by non-parametric Friedman test and Dunn’s posthoc test; p values are indicated.

We then considered if lysosome tubules display a regional pH gradient along their long axis as previously witnessed (Suresh *et al*., 2021). Here, we classified tubules as actively growing or collapsing and then measured the pHrodo/Alexa647 or Oregon green/Alexa546 fluorescence ratios along three domains of the tubule: the base (closer to cell centre), shaft, and tip (facing the cell periphery) of lysosome tubules. Interestingly, the tip of actively growing lysosomes appeared more acidic relative to the base of the tubule using pHrodo (Fig. 5D, E, Sup. Fig S5, Supplemental Videos S1, S2) or Oregon green probes (Sup. Fig. S4D-F). In comparison, the pH gradient along the axis was lost in lysosome tubules that appeared stable or collapsed (Fig. 5D, F, Sup. Fig. S4F). These results suggest that localized pH gradient at the tip of lysosome tubules may help coordinate lysosome tubulation dynamics.

### The effect of cytosolic pH on lysosome tubulation

Given the above, we postulated that the lysosomal pH generates a spatially constrained acidic aura on the cytosolic leaflet of the lysosomal membrane and that this may then modulate factors that promote tubulation. To test this hypothesis, we labelled cells with fluorescent dextran and BCECF-AM, a ratiometric cytosolic pH indicator (Bright *et al*., 1989). We generated a lysosome proximal and distal mask guided by the fluorescent dextran and measured the BCECF-AM fluorescence on those respective regions of interest in resting and LPS-treated cells (Fig. 6A, B). While the BCECF fluorescence ratio proximal and distal to lysosomes were not statistically different in resting cells, there was a statistically significant drop in this ratio proximal to lysosomes relative to cytosol distal to lysosomes in LPS-treated cells (Fig. 6A, C). This indicates that the cytosolic pH proximal to lysosomes was slightly more acidic than the cytosolic pH distal to lysosomes. We then asked if changing the cytosolic pH would impact lysosome tubulation by immersing cells in acetate Ringer’s buffer (Heuser, 1989; Durchfort *et al*., 2012) and treating cells with LPS. Indeed, lysosome tubulation was potently averted in cells incubated with acetate Ringer’s intimating that pH on the cytosolic side of lysosomes can impact the machinery that drives lysosome remodeling (Fig. 6D, E) and similar to observations with pH clamping with ionophores (Fig. 4E).

**Figure 6:**
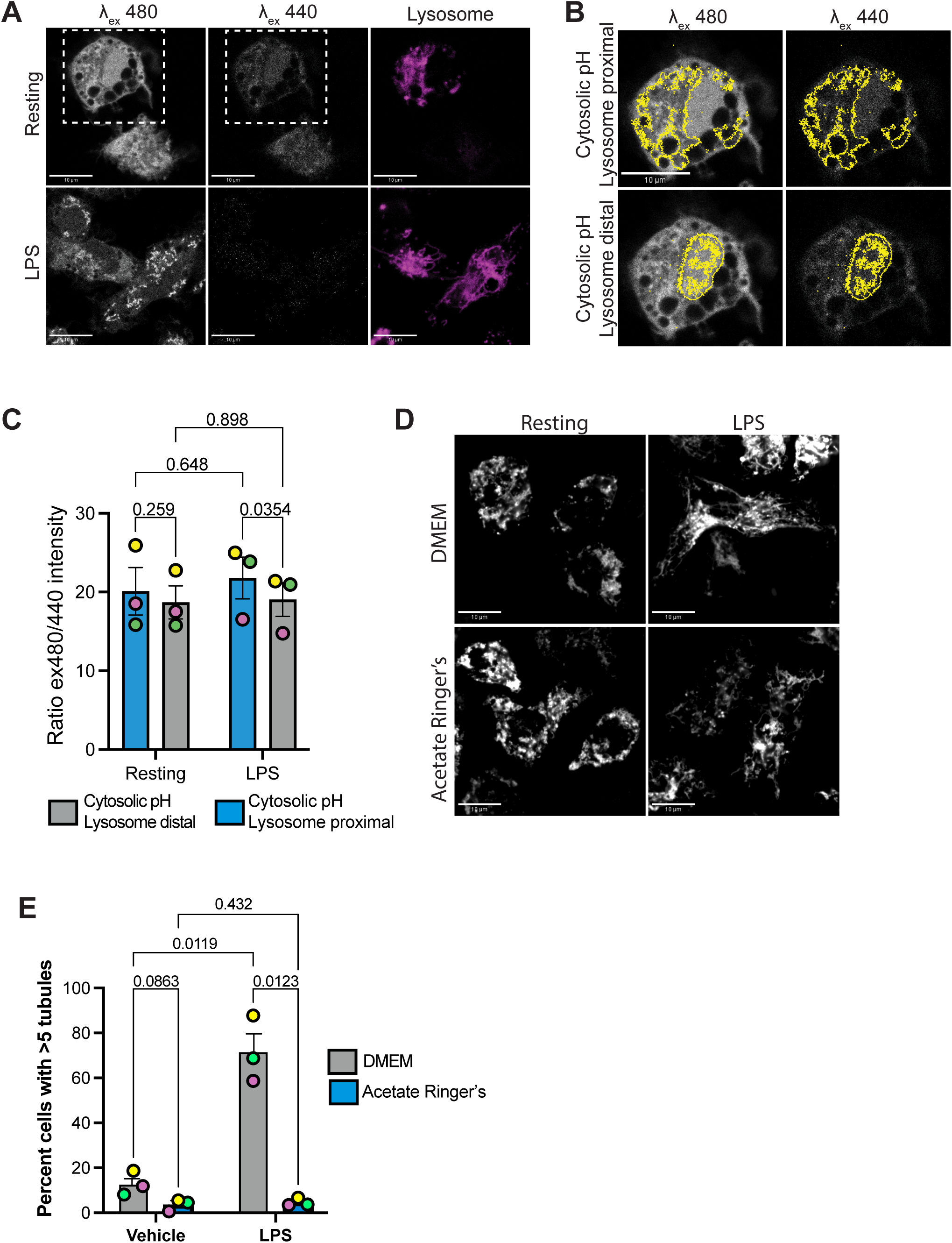
Cytosolic pH is essential for lysosome tubulation. **A.** Confocal single-plane images of live primary macrophages pre-loaded with BCECF-AM for 30 min followed by the addition of resting or LPS. Cells were excited at 440 and 480 nm, which are respectively pH insensitive and pH sensitive. **B.** Depiction of fluorescence masks using dextran and nuclear stains to form a lysosome mask and nuclear mask to estimate cytosolic pH proximal and distal to lysosomes, respectively. Representative cell is from dashed box in resting condition in A. **C.** Fluorescence ratio of BCECF-AM excited at 480 over 440 nm in resting and LPS-treated cells to estimate cytosolic pH proximal and distal to lysosomes. **D.** Single-plane confocal images of live primary macrophages with lysosomes labelled using Alexa546-dextran, followed by treatment with or without LPS in either DMEM or acetate Ringer’s medium. Scale bar = 10 µm. **E.** Cumulative percent of cells with >5 tubules. For C and E, data are shown as mean ± SEM and were analyzed by repeated measures two-way ANOVA, followed by Fisher’s LSD test.

### Lysosomal pH dissipation does not significantly disrupt microtubule organization or inhibit lysosome motility

To better understand how lysosomal pH might affect lysosome remodelling, we examined microtubule organization and lysosome motility. We compared microtubule organization by staining with α-tubulin in resting macrophages and those exposed to ConA, chloroquine, and NH_4_Cl. As a positive control for microtubule perturbation, we used nocodazole (Fig. 7A). To quantitatively determine effects on microtubule organization, we used *Skeleton* and *AnalyseSkelet*on modules in ImageJ to quantify microtubule branch number and length, and junctions (Arganda-Carreras *et al*., 2010; Schindelin *et al*., 2012). Our analysis indicates that microtubule organization remained relatively intact under these treatments, although there may be a trend towards reduced microtubule network complexity with ConA and NH_4_Cl (Fig. 7B, C). Nevertheless, this is a minor effect compared to the unequivocal loss of microtubule organization in nocodozole-treated cells (Fig. 7A-D) and the overall inhibition of lysosome tubulation (Fig. 1, 2, and 4). We thus anticipate that changes to microtubule organization is not a leading cause for loss of lysosome remodelling in cells treated with pH disruptors.

**Figure 7:**
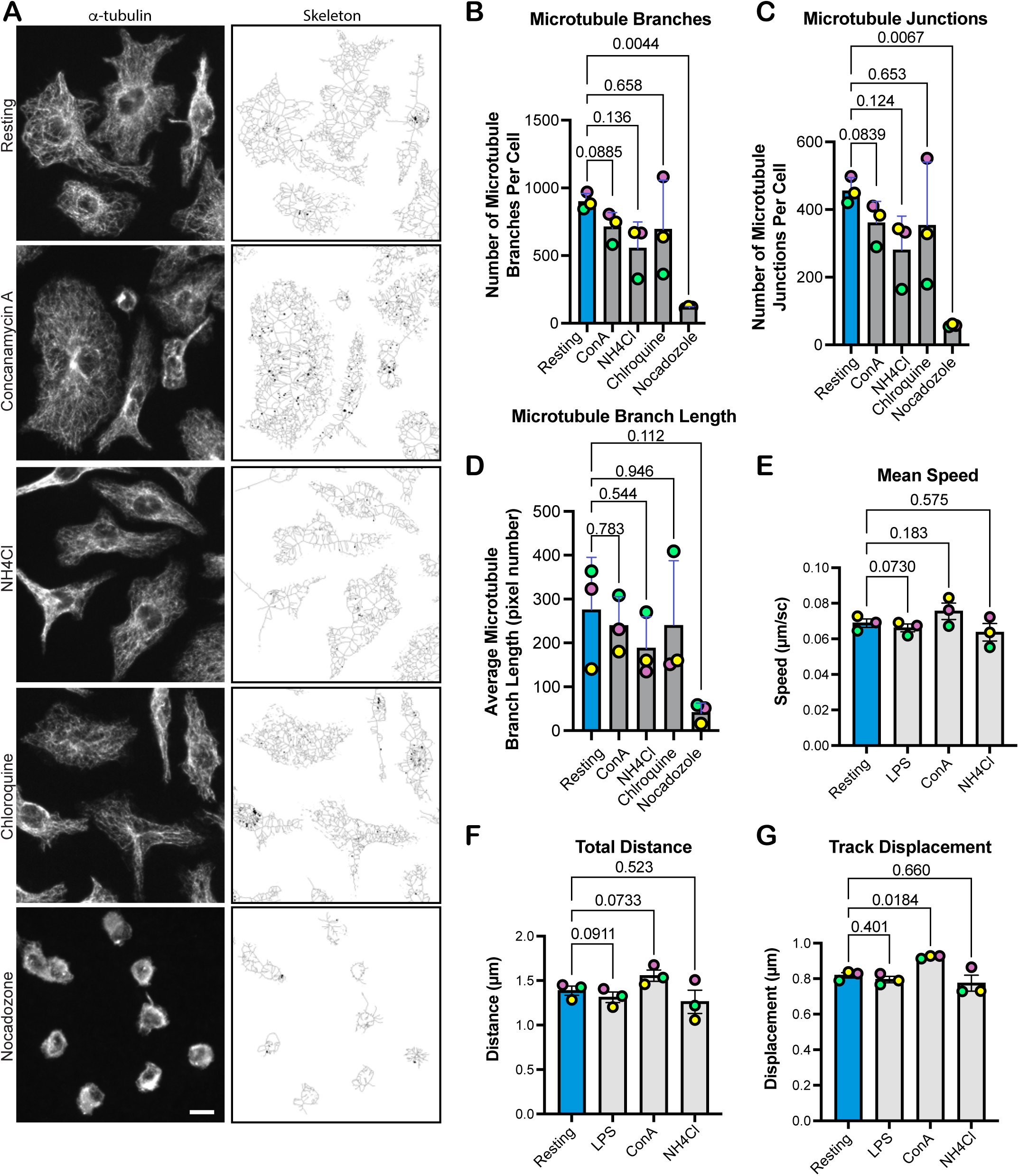
Alkalinization has relatively minor impact on microtubule organization and lysosome motility. **A.** Z-collapsed confocal images of α-tubulin-stained macrophages showing the microtubule organization. Cells were either resting, or exposed to 1 µM concanamycin A, 10 mM NH_4_Cl, 60 µM chloroquine, or 10 µM nocodazole for 1 h. Images were then converted to skeletal framework using the “Skeleton” plugin in FIJI. Scale bar = 10 µm. **B, C, D.** Skeletonized images were analysed by Skeleton 2D/3D analysis to obtain microtubule branch numbers (B), microtubule junctions (C), and microtubule branch length (D) for each condition. Data are based on 30-40 cells per condition per experiment and displayed as mean ± SEM. **E, F**, and **G**: Lysosome motility analysis showing mean speed (E), total distance (F), and track displacement (G). Data are from n=50 cells from three independent experiments and displayed as mean ± SEM. Data were tested by repeated-measures one-way ANOVA and Dunnett’s post-hoc test by comparing to resting condition; p values are indicated.

We then quantified global lysosome movement in macrophages manipulated for lysosomal pH by extracting lysosome mean speed, total distance, and displacement in cells treated with vehicle, LPS, ConA, or NH_4_Cl. Relative to resting cells, LPS treatment may slow lysosome speed and distance travelled, but our analysis is confounded by changes in lysosome morphology (Fig. 7E-G, Sup. Video S3 and S4). ConA treatment on the other hand ostensibly favours lysosome motility (Fig. 7E-G, Sup. Video S5). This was distinct from NH_4_Cl exposure, which did not significantly change any of the measured motility parameters (Fig. 7E-G, Sup. Video S6). This distinction between ConA and NH_4_Cl may reflect differences in how these compounds manipulate pH (V-ATPase inhibition vs. buffering) or distinct downstream consequences (e.g. mTORC1 inhibition). Nevertheless, we can conclude that these pH disruptors do not suppress lysosome tubulation by arresting lysosome motility, and in fact, ConA promotes motility of lysosomes. In turn, this indicates that loss of motor activity does not underpin collapse of lysosome tubules upon pH disruption.

## Discussion

Macrophages exposed to LPS remodel their lysosomes into a tubular lysosome network that may aid in antigen uptake, processing, and presentation (Hipolito *et al*., 2018; Inpanathan and Botelho, 2019). The deformation of lysosomes into tubules requires dynein and kinesin motor activity acting on a microtubule platform (Swanson *et al*., 1987b; Vyas *et al*., 2007; Mrakovic *et al*., 2012). Regulating these motors during LPS-mediated tubulation of lysosomes are the Arl8b and Rab7 GTPases and their effector motor adaptors such as FYCO1 and SKIP1 (Mrakovic *et al*., 2012). However, it is not known how these motors are productively coordinated to deform lysosome tubules in which typically the tubule grows outward towards the cell periphery (Chow *et al*., 2002; Vyas *et al*., 2007). This likely requires signaling mediators capable of forming rapid signaling niches such as ion gradients. In fact, Ca^2+^ released by TRPML1 appears to promote the ALG2-dynein adaptor and is necessary for lysosome positioning and tubulation (Li *et al*., 2016) and Ca^2+^ gradients along the tubule axis have been observed (Suresh *et al*., 2021).

Beyond Ca^2+^, lysosome pH represents another major ion modulator of lysosome function that is not only critical for lysosome degradative activity, but powers transport across the lysosomal membrane (Christensen *et al*., 2002; Mundy *et al*., 2012; Colacurcio and Nixon, 2016; Löbel *et al*., 2022; Zhang *et al*., 2022; Wu *et al*., 2023). Here, we demonstrate that the acidic lysosome pH is necessary to form and maintain lysosome tubules after LPS exposure. This role is not reflected by changes in the average cellular lysosome pH or the average pH of globular and tubular lysosomes, which did not significantly change in either resting or LPS-treated macrophages; it remains possible that this may depend on time and level of LPS stimulation (Tsang *et al*., 2000; Baker *et al*., 2015).

Strikingly, actively growing lysosome tubules display a gradient along their long axis with the peripheral end being more acidic relative to the perinuclear base. Such gradients along the long axis of lysosomes were previously observed in 50% of lysosome tubules, with the rest displaying the opposite gradient or a uniform pH along the tubule long axis (Suresh *et al*., 2021). However, here, we propose that the acidic anterior ends are associated with growing tubules. This acidic anterior tip may provide a localized niche that impacts ion flow or membrane potential that directly or indirectly governs motor activity; for example, this may stimulate kinesin-1 activity to promote growth towards the cell periphery as observed previously (Chow *et al*., 2002; Vyas *et al*., 2007). Supporting this idea, we may have detected an acidic cytosolic aura surrounding the lysosomal membrane using the BCECF-AM cytosolic pH sensor, though we could not detect pH microdomains along the lysosome length; for this, we may require more sensitive pH indicators with higher resolution such as genetically encoded pH sensors targeted to the cytosolic leaflet of lysosomes like a surface-targeted Keima or pHLare (Webb *et al*., 2021; Chen *et al*., 2025); this strategy will require generation of transgenic mice or viral transduction of these genetically encoded indicators into primary macrophages. Nevertheless, given that acetate Ringer’s and ionophores block tubulation or prevent maintenance of tubules, this indicates that pH modulates the machinery driving lysosome remodelling.

Finally, while the V-ATPase inhibition with ConA and pH alkalinization with NH_4_Cl block tubulation, they do have distinct effects. First, ConA, but not NH_4_Cl, inhibited mTORC1 activity. Second, ConA seems to promote faster lysosome motility than NH_4_Cl, despite both disrupting lysosome tubulation – thus, we propose that their negative effects on lysosome tubulation may occur through partially distinct processes that we speculate converge in loss of motor coordination. Overall, while we do not ascertain the exact function of lysosomal pH responsible for driving lysosome tubulation, we conclude that lysosomal pH is needed for both formation and maintenance of lysosome tubules in LPS-exposed primary macrophages. Moreover, we offer that differences in lysosomal pH at specific locations may help modulate cytosolic factors that sculpt globular lysosomes into tubules. Nevertheless, factors beyond pH are at play as shown by the extensive loss of lysosome tubules in chloroquine-treated cells, which appear to have vacuolated lysosomes that may result from osmotic dysregulation or altered fusion/fission dynamics. Finally, while this work was primarily driven by pharmacological manipulation of lysosome pH, it will be instructive to examine lysosome tubulation in myeloid cells derived from mice carrying activating or inhibitory mutations of V-ATPase isoforms (Zirngibl *et al*., 2019; Matsumoto *et al*., 2020; Chu *et al*., 2023; Peng *et al*., 2025).

## Materials and Methods

### Isolation, differentiation, and culture of primary macrophages

Bone marrow-derived macrophages (BMDMs) were harvested from wild-type 7-9-week-old male and female C57Bl/6 mice (Charles River Canada, Montreal, QC) as previously described with minor modifications (Inaba *et al*., 1992; Weischenfeldt and Porse, 2008). Briefly, bone marrow was isolated from femurs and tibias through perfusion with phosphate-buffered saline (PBS) using a 27-gauge syringe. Red blood cells were lysed using a hypo-osmotic treatment. Stem cells were plated according to experimental requirements in DMEM (Wisent, St. Bruno, Canada) supplemented with 10% fetal bovine serum (FBS), 20 ng/ml recombinant mouse macrophage colony-stimulating factor (PeproTech, Rocky Hill, NJ), and 5% penicillin/streptomycin antibiotics. Cells were incubated at 37 °C with 5% CO_2_. The medium was changed every two days. Experiments were conducted on days 7-8. All animals were used following institutional ethics requirements using Animal Care Protocols 209 and 461.

### Lysosome labelling for microscopy and tubulation

For lysosome labeling, BMDMs were pulsed with 100 µg/mL Alexa488-conjugated or Alexa546-conjugated dextrans, or 50-100 µg/mL Alexa647-conjugated dextran (ThermoFisher, Burlington, ON) for 30-60 min, followed by 3x wash with PBS and incubated with fresh DMEM medium for 1 h. Alternatively, cells were co-labelled with 60 μg/mL pHrodo-conjugated and 50 µg/mL Alexa 647-conjugated dextrans, or 100 µg/mL Oregon green-conjugated and 100 µg/mL Alexa546-conjugated dextrans (ThermoFisher) for 1 h to measure lysosomal pH. To induce lysosome tubulation, BMDMs were exposed to 100-500 ng/mL LPS (Invivogen, San Diego, CA) for up to 2 h.

### pH manipulation and other pharmacological treatments

To alkalinize pH, cells were incubated for 0.5 to 2 h with 1 µM concanamycin A (ConA; BioShop, Mississauga, ON) or 10 mM NH_4_Cl (Millipore-Sigma, Toronto, ON), or with 60 µM chloroquine (Millipore-Sigma, Toronto, ON) before LPS stimulation. To test the role of pH in maintaining lysosome tubulation, ConA or buffers were added after LPS-induced tubulation for 30 min. To clamp the cytoplasm and lysosomes to specific pH, we treated cells with 2 µM monensin and 5 µM nigericin in sodium rich-medium (140 mM NaCl, 5 mM glucose, and 15 mM HEPES) set to pH 7.4, 5.0, or 4.0 using KOH or HCl for 10 min and then imaged immediately. To acidify the cytosol, cells were incubated with acetate Ringer’s medium for 30 min followed by LPS to induce tubulation. Other drugs used included 100 nM torin1 or 10 µM nocodazole for 1 h.

### Immunofluorescence

BMDMs were fixed using 4% paraformaldehyde (PFA) for 15 min after specific treatments. Cells were washed with PBS three times, then treated with permeabilization buffer (0.1% Triton-X in PBS), and blocked for 1 h with 3% BSA. Cells were then incubated with 1:200 rabbit anti-α-tubulin antibodies (Sigma-Aldrich, St. Louis, MO). After washing excess primary antibodies, cells were incubated with Alexa-dye conjugated donkey polyclonal antibodies against rabbit immunoglobulin (1:1000; ThermoFisher) for 1 h. Nuclei were stained with 0.4 µg/mL DAPI for 5 min. Cells were then mounted on a slide using DAKO mounting medium, and dried for 24 h before imaging/storage at 4 °C.

### Fluorescence microscopy

All live-cell imaging was done in 5% CO_2_ and at 37 C using an environmental control chamber. Fixed and the majority of live-cell imaging was done with either a *i)* Quorum Diskovery Spinning Disc Confocal Microscopy system equipped with a Leica DMi8 inverted microscope connected to an Andor Zyla 4.2 Megapixel sCMOS camera and a iXon 897 EMCCD camera, controlled by Quorum WaveFX powered by MetaMorph software (Quorum Technologies, Guelph, ON); or *ii)* an Olympus Spinning Disk Confocal Microscope system equipped with a IX83 P2ZF microscope connected to a ORCA-Flash4.0 V3 Digital CMOS camera powered by the CellSens Dimension software workflow (Evident Scientific, Tokyo, Japan). For lysosome motility measurements, cells were imaged with the Re-scan confocal microscope system mounted on a Leica DMi8 inverted microscope, equipped with an ORCA Flash4.0 V3 sCMOS camera, and controlled by Volocity Software (Quorum Technologies). Images were acquired at 0.25 hz for 1 min. Live-cell movies to measure lysosome pH of lysosome tubules, images were acquired at a rate of one frame for every 500 ms for a total duration of 3 min. The dual mode acquisition mode was utilized wherein images were acquired in both 488 and 546/647 channels at the same time.

### Analysis of microtubule structure

Single-plane images were converted to 8-bit images using FIJI/ImageJ followed by application of fluorescence intensity threshold to select microtubules. Images were converted to binary and filaments analysed through *Skeleton* and *AnalyzeSkeleton* plugin (Arganda-Carreras *et al*., 2010; Schindelin *et al*., 2012). Total number of microtubules junctions, where junctions represent filamentous pixels from where two or more microtubule branches arise, total number of microtubule branches and average microtubule branch length were scored and collected for data analysis.

### Analysis of lysosome tubulation and pH

For manual quantification of lysosome tubules, lysosomes were considered tubular if their length was over 4 µm and apparent branches were counted as a separate tubule. To estimate relative lysosome pH, pH-sensitive channel (pHRodo or Oregon green) and pH-insensitive channel (Alexa647 or Alexa546) were co-acquired. Images were background corrected and then assessed for fluorescence intensity for both channels. The global lysosomal pH was estimated by calculating the ratio of both channels for the whole cell. To estimate pH of puncta and tubular lysosomes individually, we employed circularity and Feret’s diameter/length ratio: lysosomes were defined as globular if circularity was >0.5 and the Feret’s ratio was <1.9; lysosomes were defined as tubular if circularity was <0.5 and Feret’s >1.9. = puncta). For pH analysis along the long axis of lysosome tubules, regions of interest (ROIs) were drawn manually in FIJI based on Alexa647/Alexa546 channel and then ratiometric values were extracted for each region: base is defined as perinuclear/central end; shaft as the middle of the tubule, and tip as the end towards the cell periphery.

### Lysosome labelling for flow cytometric pH measurements

For flow cytometric pH measurements, BMDMs were loaded with 50 µg/ml AlexaFluor 546-conjugated dextran, and either 50 µg/ml Oregon Green 488-conjugated dextran, or 50 µg/ml pHrodo Green-conjugated dextran for 1 h, followed by 3x wash with PBS and incubated with fresh DMEM, 10% FBS, for 30 min. To induce lysosome tubulation, BMDMs were exposed to 100 ng/ml LPS for 2 h prior to analysis. As a control, separate samples of BMDMs were treated with either 60 µM chloroquine (Cell Signalling Technology) or 10mM NH_4_Cl (Millipore-Sigma) for 30 minutes prior to analysis. Following treatment, cells were washed 3x with PBS and gently lifted into phenol red-free DMEM, 10% FBS, 25 mM HEPES containing the respective treatment to maintain live, active lysosomes. A Beckman Coulter Cytoflex S analyzer was used to measure the mean fluorescence intensity in both the PE (AlexaFluor 546-conjugated dextran) and FITC (Oregon Green 488-and pHrodo Green-conjugated dextrans) channels of approximately 10,000 cells per sample.

### BCECF-AM ratiometric imaging for cytosolic pH

To measure the relative cytosolic pH, cells with Alexa647 dextran-loaded lysosomes were treated with 1 μM 2’,7’-Bis-(2-Carboxyethyl)-5-(and-6)-Carboxyfluorescein-Acetoxymethyl Ester (BCECF-AM; Invitrogen, Carlsbad, CA) in Hanks’ Balanced Salt Solution (HBSS; ThermoFisher) for 30 min. After washing excess BCECF-AM, cells were treated for 0.5 h to 2 h with DMEM or 500 ng/mL LPS in DMEM. Nuclei were stained with 200 µg/mL Hoechst (Bio-Rad Laboratories) for 10 min. Live cell imaging was then performed on the Leica Stellaris 8 confocal microscope mounted on a Leica DMi8 inverted microscope and equipped with hybrid (HyD) detectors powered by the Leica Application Suite X (LAS X) software. Excitation was done at 440 nm (pH insensitive) and 480 nm (pH sensitive) and emission captured at ∼535 nm Single-plane images were converted to 8-bit images using FIJI followed by application of fluorescence intensity threshold to select lysosomes and nuclei separately, creating a mask of lysosomes from the dextran channel (lysosome proximal cytosolic pH) and a mask of the nuclei from the Hoechst channel (lysosome distal cytosolic pH). The ratio of fluorescence intensity at 480 nm to 440 nm (Ex480/Ex440) was used as a measure of relative cytosolic pH, with higher ratios corresponding to a more alkaline pH and lower ratios corresponding to a more acidic pH.

### Analysis of lysosome motility

Time-lapse images of lysosomes were analysed for lysosome speed, distance travelled, and displacement over 1 min using FIJI TrackMate plugin (Tinevez *et al*., 2017; Ershov *et al*., 2022). We used standard settings, and the LoG detector with a blob diameter and threshold set to 3 pixels in diameter, followed by the Simple LAP tracker, linking and gap closing distance were also set to 3 pixels to minimize “jumping”. This allowed about 40-60% of lysosomes to be tracked. Tubular lysosomes were tracked by virtue of puncta assembled along their length as this program does not allow one to track entire tubules.

### Training of machine learning model

Single-plane TIFF images obtained from the spinning disk microscope were converted to 8-bit files using ImageJ software. Single macrophages were cropped out to size 300×300 pixels and saved as JPG files.

A convolutional neural network based on ResNet50 pre-trained on ImageNet was used to classify cell images into two classes: macrophages with tubulating or non-tubulating lysosomes. The model was implemented in Python using Keras API with TensorFlow back-end, and all analyses were performed in Jupyter notebooks. The ResNet50 model was configured to accept inputs of size 300 × 300 × 3, and all images were resized accordingly. Pixel intensities were scaled to the [0, 1] range by dividing by 255. For transfer learning, 140 images of non-tubulating lysosomes and 199 images of tubulating lysosomes were used to fine-tune the model. The dataset was split into training and test subsets using an 80-20 ratio. Data augmentation, including rotations, flips, translations, and brightness adjustments, was applied to the training images to mimic experimental variability and improve generalization. ResNet50’s global average pooling layer was retained to reduce feature dimensionality before the classification layer. A Dense layer with two output units and softmax activation was added to perform binary classification. The model was compiled using the Adam optimizer with a learning rate of 0.0001 and trained with categorical cross-entropy loss. The lower learning rate was chosen to ensure stable fine-tuning of the pre-trained ResNet50 weights with our small training set. Training was performed for up to 25 epochs with a batch size of 5, and early stopping was applied based on validation accuracy to prevent overfitting. The trained model was then used to classify new images. For each image, the model output probabilities for both classes, and the class with the higher probability was assigned as the predicted label. Accuracy and confusion matrices were calculated to assess model performance.

To score lysosome tubulation, we used the model-generated probability that a given cell belonged to the “cells with lysosome tubules” category, expressed as a percentage. The mean probability of 50 cells per experimental group per experiment was calculated for each treatment and compared to the resting condition to evaluate changes in lysosome tubulation upon treatment.

### SDS-PAGE and Western blotting

After specific treatments, cells were lysed with 2x Laemmli buffer supplemented with 1:100 protease inhibitor cocktail (Sigma-Aldrich) and PhosSTOP protease inhibitor (Roche, Mississauga, ON). Proteins were then separated in a 10% SDS-PAGE, followed by protein transfer to a polyvinylidene difluoride (PVDF) membrane (EMD Millipore) and blocked in 1% BSA in Tris-buffered saline buffer with 0.1% Tween-20 (TBST). Membranes were then immunoblotted using primary and secondary antibodies prepared in 5% BSA in TBST at the dilutions indicated. Primary antibodies used were rabbit anti-p70 S6 kinase, phosphoThr389-p70 S6K, and β-actin (Cell Signalling Technologies, Danvers, MA) all at 1:1,000. We used secondary HRP-linked antibodies raised in donkey (1:10,000, Bethyl). Proteins were detected using Clarity Enhanced Chemiluminescence (Bio-Rad Laboratories, Mississauga, ON) with a ChemiDoc Touch imaging system (Bio-Rad). Protein quantification was performed using Image Lab software (Bio-Rad), where protein loading was normalized to levels of b-actin or total S6K signal.

### Statistical analyses

All experiments were completed at least three independent times. For microscopy analysis, 15-300 cells were quantified per experiment per condition. For individual lysosome analysis, 50-100 lysosomes were analysed. When selecting individual lysosomes, we used a pre-determined grid to select these and reduce bias. Flow cytometry used 10,000 cells per experiment per condition. Data were typically represented as a mean ± standard error of the mean (SM) unless otherwise stated in figure legends. Data were assumed to be normally distributed, unless normalized to 1. Depending on these assumptions, we used Student’s t-test (two conditions, non-normalized data) or Wilcoxon one-sample t-test (normalized data, two conditions), repeated measures one-way ANOVA (three or more conditions), repeated measures two-way ANOVA with matched data (two parameters with two or more conditions), or Friedman test (if data is not normally distributed with three of more conditions). Where applicable, Geisser-Greenhouse correction was applied as recommended by GraphPad Prism. Post-hoc tests were used as recommended by GraphPad Prism and are indicated in the figure legends. A p>0.05 was set as the significance cut off, but we display the actual p values for full disclosure, noting that p-values above but near p=0.05 may indicate a possible biological effect.

## Abbreviations

BMDM: bone marrow-derived macrophages;
ConA: concanamycin A;
LPS: lipopolysaccharide;
SEM: standard error of the mean

## Funding and acknowledgements

We would like to thank the staff at the Vivarium facilities in St. Michael’s Hospital and Unity Health Toronto for their support and guidance in working with mice. This work was funded by grants from the Canadian Institutes of Health Research (PJT-166047, PJT-197934), the Canada Fountain for Innovation (32957, 38151), and the Canada Research Chairs Program (950-232333) with contributions from Toronto Metropolitan University and Ontario Ministry of Economic Development, Job Creation, and Trade (38151) to RJB. Work in MMU’s lab is supported by a Discovery Grant from the Natural Sciences and Engineering Research Council of Canada (NSERC-DG 03741-2021).

## Supplemental Materials

**Supplemental Figure S1: Validation of concanamycin A deacidifcation of lysosomes in macrophages. A.** Confocal images of macrophages co-labelled with Alexa647-and pHrodo-dextrans in resting and LPS-treated states with and without concanamycin A. Scale bar = 10 µm. **B.** The mean ± SEM of pHrodo/Alexa647 fluorescence ratio from indicated conditions from n=3 independent experiments with 30-40 cells per condition per experiment. Data were analysed with repeated-measures two-way ANOVA with Tukey’s post-hoc test; p values are shown.

**Supplemental Figure S2: Effect of NH_4_Cl and chloroquine on lysosome pH in macrophages. A.** Confocal images of macrophages co-labelled with Alexa647-(magenta in merge) and pHrodo-(green in merge) dextrans in resting, concanamycin A, chloroquine, and NH_4_Cl treatment. Scale bar = 10 µm. **B.** The mean ± SEM of pHrodo/Alexa647 fluorescence ratio from indicated conditions and from n=3 independent experiments with 25-30 cells per condition per experiment. **C.** The mean ± SEM of pHrodo/Alexa647 fluorescence ratio per cell measured by flow cytometry from N=3 independent experiments and 10,000 cells per condition per experiment. **D.** The mean ± SEM of Oregon green/Alexa647 fluorescence ratio per cell measured by flow cytometry from N=3 independent experiments and 10,000 cells per condition per experiment. For C and D, cells were resting or treated with LPS, or ConA, or NH_4_Cl. For B, C, and D, data were analysed with repeated-measures one-way ANOVA with Dunnett’s post-hoc test comparing to resting conditions; p values are shown.

**Supplemental Figure S3: Machine-learning model for lysosome tubulation. A.** Shown are three representative cells for the two manually assigned classes of cells: cells with non-tubular lysosomes and cells with tubular lysosomes. **B.** Example of validation image set previously scored manually as non-tubular [0, x] or tubular [1, x] and then scored by the trained model as non-tubular [x, 0] or tubular [x, 1]. Agreement existed when images receive a [0,0] or [1,1]. **C.** The binary supervised machine-learning classifier achieved > 80% accuracy

**Supplemental Figure S4: Oregon green-labelled lysosomes indicate that LPS does not change the global lysosomal pH, but growing tubules are more acidic at their anterior tips. A.** Confocal images of resting and LPS-treated primary macrophages labelled co-labelled with Oregon green dextran (pH sensitive) and Alexa546-dextran (pH insensitive); merge shows two channels (grayscale = Oregon green; magenta = Alexa546). Scale bar = 10 µm. **B**. Average cellular Oregon green to Alexa546 fluorescence intensity in resting and LPS-treated macrophages. **C**. Mean Oregon green to Alexa546 fluorescence intensity of punctate and tubular lysosomes in resting and LPS-treated macrophages. **D.** Timelapse of ratiometric images of Oregon green/Alexa546 fluorescence of LPS-treated macrophages. The ratiometric images are pseudocoloured corresponding to pH shown in the LUT. The purple arrow identifies a tubular lysosome peripheral tip growing and showing more acidic ratio relative to its base. Time is in seconds. **E, F.** Normalized mean ratio of Oregon green/Alexa546 fluorescence along the long axis of tubules observed to grow (E) or unchanged/collapsing (F); regions along the tubule axis were defined as the base (beginning of tubule at the perinuclear region), shaft (centre of the tubule), and tip (tubule end towards the cell periphery). All data are means ± SEM from at least three independent experiments. Data in B were from n=50 cells per experiment per condition and were tested by a paired, two-tail Student’s t-test; Data in C were based on n=25 cells and 1800-2000 individual lysosomes per experiment per condition and was tested with a repeated measures two-way ANOVA and Tukey’s post-hoc test. Data in E and F were assessed by repeated measures one-way ANOVA and Tukey’s post-hoc test; p values are indicated.

**Supplemental Figure S5: Additional example of pH gradient along the long axis of growing lysosome tubules.** Timelapse of ratiometric images of pHRodo/Alexa647 fluorescence of LPS-treated macrophages. The ratiometric images are pseudocoloured corresponding to pH shown in the LUT. Purple arrow identifies a tubular lysosome peripheral tip growing and showing more acidic ratio.

**Supplemental Video 1: Ratiometric image of lysosomal pH in LPS-treated macrophage** showing a short tubule collapsing and another one extending. See Sup. Fig. S4 or additional description and pseudocolour information. Time in seconds.

**Supplemental Video 2: Ratiometric image of lysosomal pH in LPS-treated macrophage** showing a short tubule collapsing and another one extending. See Sup. Fig. S4 for additional description and pseudocolour information. Time in seconds.

**Supplemental Video 3: Lysosome motility of resting macrophages.** Live-cell confocal imaging of Alexa546-dextran-labelled macrophages at 10 sec/frame for 10 min.

**Supplemental Video 4: Lysosome motility of LPS-treated macrophages.** Live-cell confocal imaging of Alexa546-dextran-labelled macrophages at 10 sec/frame for 10 min.

**Supplemental Video 5: Lysosome motility of ConA-treated macrophages.** Live-cell confocal imaging of Alexa546-dextran-labelled macrophages at 10 sec/frame for 10 min.

**Supplemental Video 6: Lysosome motility of NH_4_Cl-treated macrophages.** Live-cell confocal imaging of Alexa546-dextran-labelled macrophages at 10 sec/frame for 10 min.

## Notes

### Competing Interest Statement

The authors have declared no competing interest.

### Summary of Updates

Minor changes to text and correction to author's name Berilia to Beriia.

