## Supplementary figures and images for "Dissipation of lysosome pH impairs formation and collapses existing LPS-induced lysosome tubules in macrophages"

### Supplemental Figure S1

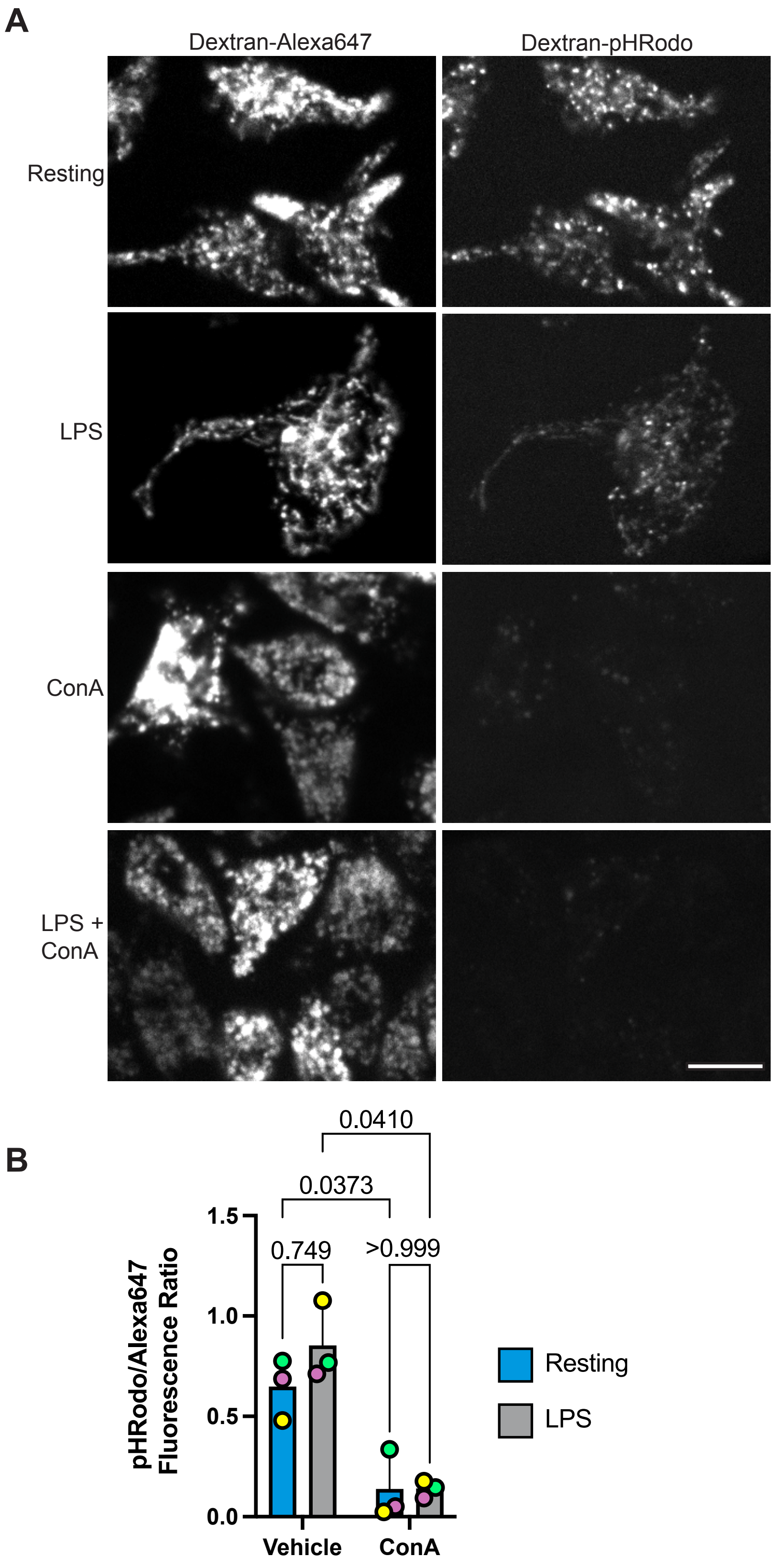


Supplemental Figure S1

### Supplemental Figure S2

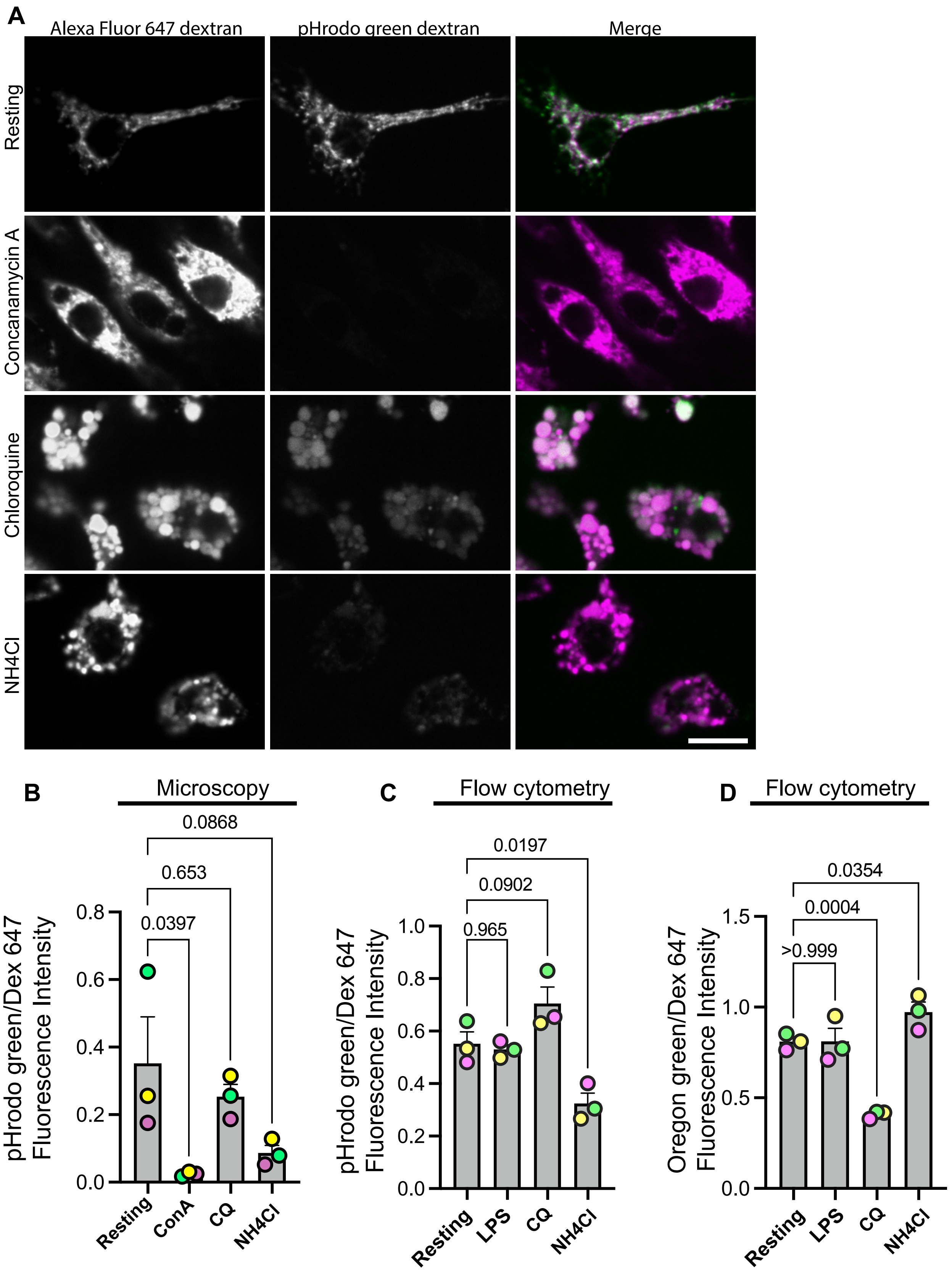


Supplemental Figure S2

### Supplemental Figure S3

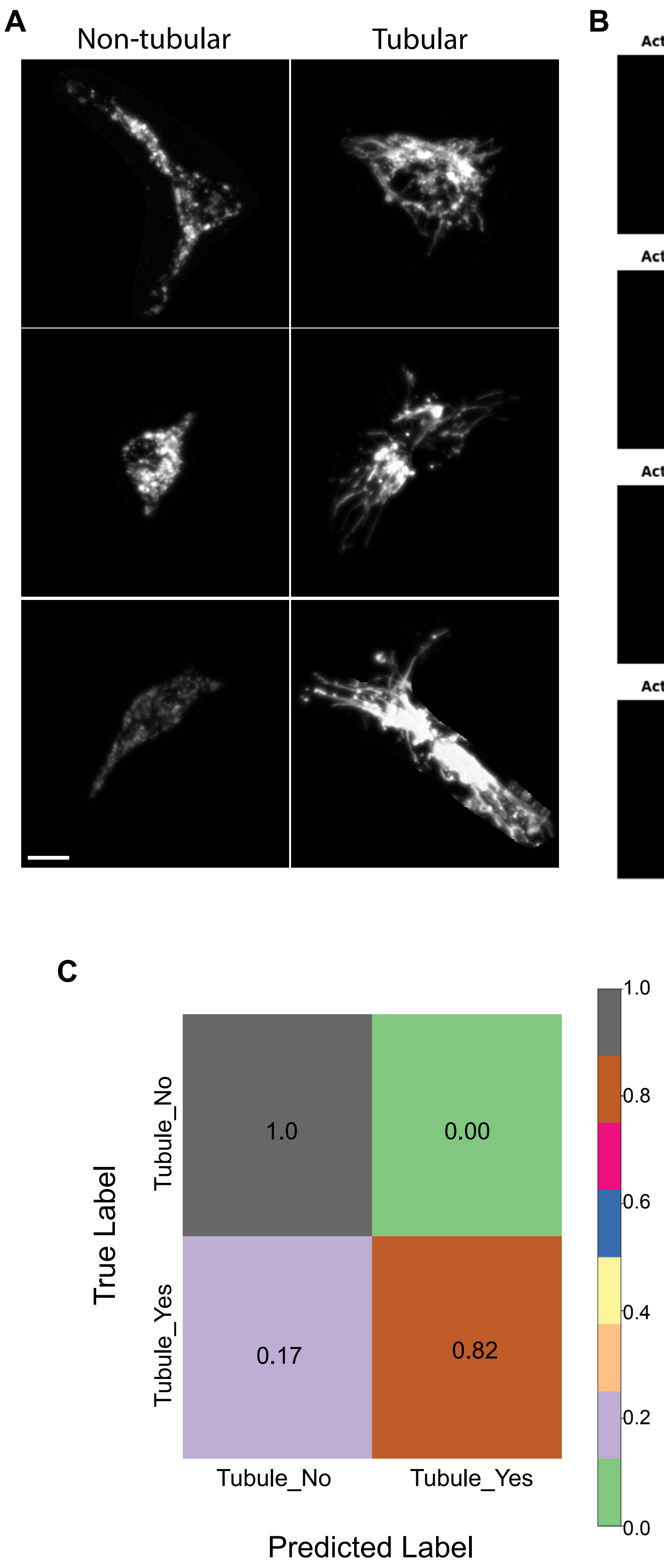


Supplemental Fig. S3

### Supplemental Figure S4

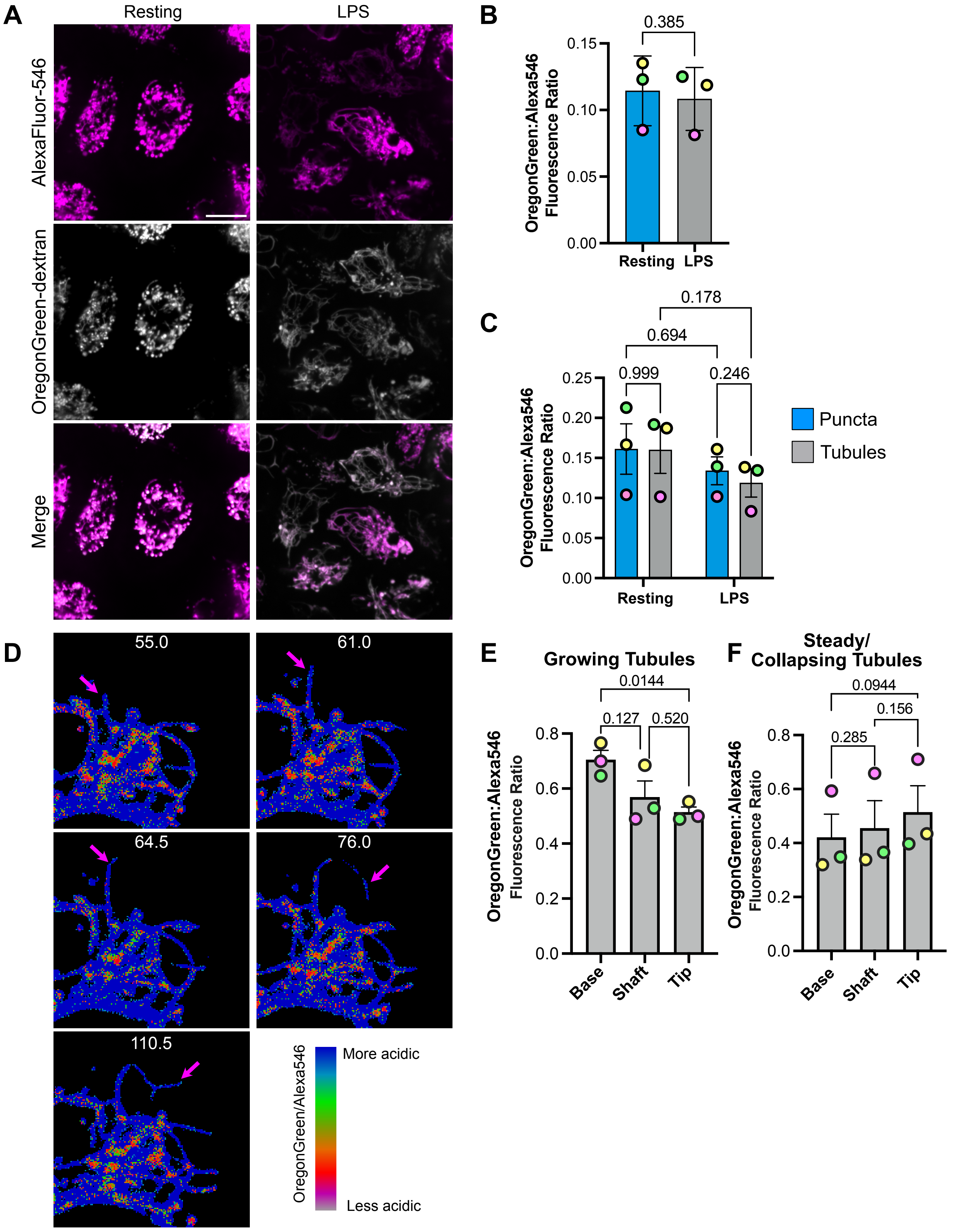


**Supplemental Figure S4**

### Supplemental Figure S5

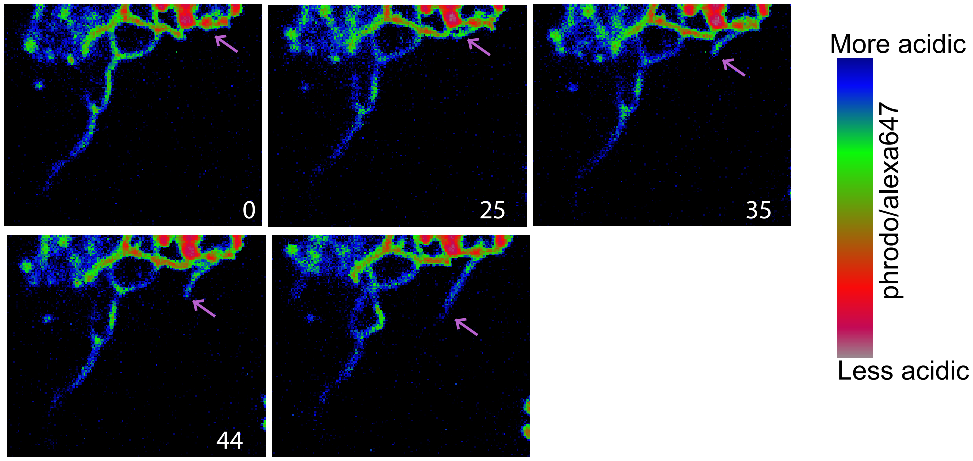


Supplemental Figure S5
